# Cyanamide-inducible expression of homing nuclease ^I-^*Sce*I for iterative genome engineering and parallel promoter characterisation in *Saccharomyces cerevisiae*

**DOI:** 10.1101/2024.03.26.586389

**Authors:** Liam McDonnell, Samuel Evans, Zeyu Lu, Mitch Suchoronczak, Jonah Leighton, Eugene Ordeniza, Blake Ritchie, Nik Valado, Niamh Walsh, James Antoney, Chengqiang Wang, Colin Scott, Robert Speight, Claudia E. Vickers, Bingyin Peng

**Affiliations:** Centre of Agriculture and the Bioeconomy, School of Biology and Environmental Science, Faculty of Science, Queensland University of Technology, Brisbane, QLD, 4000, Australia; ARC Centre of Excellence in Synthetic Biology, Australia; Australian Institute for Bioengineering and Nanotechnology (AIBN), The University of Queensland, Brisbane, QLD, 4072, Australia; School of Biology and Environmental Science, Faculty of Science, Queensland University of Technology, Brisbane, QLD, 4000, Australia; College of Life Sciences, Shandong Agricultural University, Taian, Shandong Province, 271018, People’s Republic of China; CSIRO Environment, Black Mountain Science and Innovation Park, Canberra, ACT 2601, Australia; Advanced Engineering Biology Future Science Platform, Commonwealth Scientific and Industrial Research Organisation (CSIRO), Black Mountain, ACT, 2601, Australia

## Abstract

In synthetic biology, microbial chasses including yeast *Saccharomyces cerevisiae* are iteratively engineered with increasing complexity and scale. Wet-lab genetic engineering tools are developed and optimised to facilitate strain construction but are often incompatible with each other due to shared regulatory elements, such as the galactose-inducible (*GAL*) promoter in *S. cerevisiae*. Here, we prototyped the cyanamide-induced ^I-^*Sce*I-mediated double-strand DNA breaks (DSBs) for selectable marker recycling in yeast metabolic engineering. We further combined cyanamide-induced ^I-^*Sce*I-mediated DSB and maltose-induced MazF-mediated negative selection for plasmid-free *in situ* promoter replacement, which simplified the molecular cloning procedure for promoter characterisation in *S. cerevisiae*. We then characterised three tetracycline-inducible promoters of differential strength, a non-leaky β-estradiol-inducible promoter, cyanamide-inducible *DDI2* promoter, bidirectional *MAL32/MAL31* promoters, and five pairs of bidirectional *GAL1/GAL10* promoters. Overall, alternative regulatory controls for genome engineering tools are important for the construction of complexed genotypes in microbial systems for synthetic biology and metabolic engineering applications.

## Introduction

In synthetic biology and metabolic engineering, yeasts have been intensively engineered to produce and improve production of certain fuels, chemicals, and other bioproducts[1-4]. It often requires genetic modification on multiple genes and at multiple genomic loci to generate ideal genotypes and phenotypes[5, 6]. Such modification capacity is rapidly evolving as the result of developing various engineering tools, including computational design/learn tools, laboratory build/test tools, and high throughput procedures assisted by robotic automation [7-11]. Wet-lab strain engineering is essentially important in this process but often labour-intensive and resource-consuming [12]. To accelerate the process, strain engineering tools and strategies can be further explored.

Genomic integration of transgenes in the common yeast *Saccharomyces cerevisiae* was first reported in the 1970s[13]. Following this, genomic engineering methodologies and strategies, including plasmid vectors[14, 15], positive/negative selectable markers[16-18], multi-gene introduction[19-21], and large-scale genome synthesis[22, 23], have been continuously developed and improved. In rational yeast engineering, gene knock-out and gene knock-in are essential to deliver identical genotypes according to the research design (or hypotheses). Inherit advantages in *S. cerevisiae* include its highly efficient homologous recombination, whereas other non-conventional yeasts, like *Komagataella phaffii* (previously known as *Pichia pastoris*), may be engineered to reach the comparable efficiency as well[24, 25]. Through homologous recombination, endogenous genes can be deleted through a knock-out strategy and heterologous genes can be integrated through a knock-in strategy. To facilitate knock-out/knock-in engineering processes, molecular biology tools are deployed, i.e., tyrosine recombinase-mediated site-specific recombination (like Cre and Flp)[26], CRISPR-associated nuclease-mediated genome double-strand breaking (like Cas9 and Cbf1)[27], meganuclease-mediated genome double-strand breaking (like the homing nuclease I-*Sce*I)[28], and serine recombinase-mediated site-specific recombination (like Bxb1 and PhiC31)[29]. These tools may be further modified for alternative purposes. For example, Cre-loxP recombination is used for SCRaMbLE[30], synthetic chromosome rearrangement and modification by loxP-mediated evolution. CRISPR tools are used for CRISPRi (CRISPR interference) [31] and CRISPRa (CRISPR activation)[32]. Ideally, all these tools can be compatible with each other to realise their functions in a single host, allowing the engineering and integration of complexed biological systems.

Some of these molecular biology tools are not compatible with each other for parallel applications in the host. For example, galactose-inducible promoters are used to control the induction of Cre-LoxP-mediated SCraMbLE[33], Cre-LoxP-mediated marker removal[34], ^I-^*Sce*I-mediated marker excision[28], ^I-^*Sce*I-mediated on-genome assembly of multiple genes[19], and mostly the induction of heterologous metabolic pathways in metabolic engineering[35-37]. To solve the compatibility problem, orthogonal genetic regulatory systems can be used to control each of these engineering machineries. In *S. cerevisiae*, both native and synthetic regulatory systems have been widely investigated in synthetic biology and yeast engineering studies[3, 38, 39]. The cyanamide-inducible promoter (*DDI2*) and maltose-inducible promoter (*MAL32*) are the options of native systems, and do not require the introduction of additional synthetic regulatory factors [40-42]. Therefore, we aimed to explore their usage for the orthogonal control on genome engineering tools.

In this study, we focused on exploring the ^I-^*Sce*I-mediated genome engineering by using cyanamide-inducible *DDI2* promoter to control ^I-^*Sce*I expression. In one application, a cyanamide-inducible ^I-^*Sce*I system was used for the removal of the selectable marker. In another application, this system was used to facilitate the knock-in of a yeast promoter upstream of the fluorescent protein reporter gene(s), whereas maltose-inducible expression of the bacterial toxin MazF [18] was deployed to facilitate the selection. This tool allows for the characterisation of multiple yeast promoters in parallel and independently from plasmid cloning.

## Materials and Methods

### Plasmid and strain construction

Yeast strains, plasmids, primers used in yeast colony PCR, and the *CYC1core[4×Z268+1×LmrO]* promoter sequence are listed in Supplementary Data File 1. Plasmid sequences are in Supplementary Data File 2.

### Yeast cultivation and transformation

Yeast extract Peptone media (YP media; 10 g L^-1^ yeast extract, and 20 g L^-1^ peptone), supplemented with 20 g L^-1^ glucose (YP-glucose media), were used in general strain maintenance and development. Antibiotic G418 sulphate (100 μg mL^-1^) or (300 μg mL^-1^) hygromycin B was used in the selection of the strain harbouring the *KanMX4* selectable marker[34]. Yeast nitrogen base media (YNB media; 6.9 g L^-1^ yeast nitrogen base without ammonia sulphate and ammino acids, pH 6.0), were used to grow the strain harbouring the *URA3* selectable marker in yeast transformation and characterisation. When maltose and/or galactose were used as the carbon source(s), each of them was supplemented at the concentration of 20 g L^-1^. For characterisation of genetic induction, 2 mM cyanamide (diluted from 4 M stock in H_2_O), 125 μM tetracycline (diluted from 125 mM stock in dimethylsulfoxide:ethanol 1:1 mixture; stored at -80 °C), or 1 μM β-estrodiol (diluted from 1 mM stock in dimethylsulfoxide) was used. Agar (20 g L^-1^) was added to prepare the solid media.

*S. cerevisiae* transformation was performed using the LiAc/ssDNA/PEG method with modifications[43]. Yeast cells from the recovery plate were inoculated into YP-glucose media at OD600 = 0.001 for routine transformation or 0.0004 for the transformation with two-hour cyanamide induction and grown overnight. The LiAc/SScarrierDNA/PEG method [43] was followed to prepare the transformation mixture with heat-shock treatment. For antibiotic selection, cells collected from transformation mixture were precultured for at least three hours in YP-glucose media and then spread onto the selection agar plates for further cultivation. For the yeast promoter knock-in with the negative selection through maltose-induced MazF toxin expression, cells were precultured in 1 mL YP-glucose media and incubated at room temperature overnight, and 10 μL of overnight culture was spread onto each YP-maltose agar plate supplemented with the required inducer like galactose, tetracycline, cyanamide, or β-estrodiol.

For yeast promoter characterisation, yeast cells were pre-cultured in 200 μL YNB-glucose media in 5 ml sterile falcon tubes for ∼24 hours to the stationary phase. The cultures were grown in a 200 rpm 30 °C incubator. For characterisation in YNB-glucose media, the precultures were diluted by 1000 times in 200 μL YNB-glucose media with or without inducer supplement. For the characterisation in YNB-galactose media, the precultures were diluted by 100 times in 200 μL YNB-galactose media. For the characterisation in YP media, the precultures were diluted by 10000 times in 200 μL fresh YP-glucose or YP-maltose media. The diluted cultures were grown overnight for fluorescence measurements in single cells through flow cytometer.

### Yeast colony PCR

Yeast colony PCR were performed using the protocol we published previously[5]. Yeast cells were resuspended in 5 μL of yeast cell digestion solution, the 1:10 mixture of 1 × phosphate-buffered saline (PBS) buffer and Zymolyase-20T (nacalai tesque, Japan) stock solution (1U per μL in 1*PBS buffer containing 100 mM DTT and 50% v/v glycerol). Yeast cells were digested at 37 °C for 30 minutes, denatured at 95 °C for 5 min, and diluted with 100 μL of water. The mixture (1 μL) was used as the template in 10 μL of PrimeSTAR GXL DNA Polymerase (Takara Bio Inc., Japan) reaction. The typical PCR thermal-cycle conditions are (1) 98 °C for 30 s, (98 °C for 15 s, 50 °C for 10 s, 68 °C for 5 min)*3, (98 °C for 15 s, 55 °C for 10 s, 68 °C for 5 min)*27, 68 °C for 5 min. The extension at 68 °C was adjusted according to the amplicon length.

### Flow cytometry

A BD FACSCanto II flow cytometry (Firmware version 1.49) was used to measure yEGFP [44] and E2Crimson [45] fluorescence in single cells. Yeast cells in overnight cultures were used directly for fluorescence analysis. yEGFP fluorescence was monitored at the FITC-A channel with a 488 nm laser. E2Crimson fluorescence was monitored at the APC-A channel with a 633 nm laser. Mean values of FSC-A (PMT voltage = 200; gating threshold = 3000), SSC-A (PMT voltage = 200), FITC-A (PMT voltage = 450), and APC-A (PMT voltage = 535) for all detected events were extracted using BD FACSDiva Software (version 8.0.1; CST Version 3.0.1; PLA version 2.0). yEGFP and E2Crimson fluorescence levels were expressed as the percentage of the average background autofluorescence from the exponential-phase cells of FP-negative reference strain CEN.PK113-7D[46].

## Results

### Cyanamide-inducible ^I-^*Sce*I expression for selectable marker excision in iterative strain engineering

Galactose-inducible *GAL1* promoter has previously been used to control the expression of ^I-^*Sce*I[19, 28]. ^I-^*Sce*I recognises and causes the double-strand break (DSB) at the 18 bp ^I-^*Sce*I recognition site (5′ -AGTTACGCTAGGGATAACAGGGTAATATAG-3 ′). DSBs trigger DNA repair, prevalently through homologous recombination in *S. cerevisiae* when homologous sequences surround ^I-^*Sce*I recognition site. This system has demonstrated very efficient for selectable marker excision[28]. We have previously used galactose-inducible *GAL* promoters to control the expression of heterologous terpene synthetic pathways[5, 14, 47]. Induction of terpene synthesis may cause metabolic burden, observed as the slow growth phenotype. To circumvent the metabolic burden during strain development and maintenance, we developed a synthetic genetic circuit for tetracycline-mediated repression of *GAL* promoters [38]. We aimed to use alternative regulatory promoters to control genome-editing tools. The *DDI2* promoter was first considered because of its binary ‘OFF-to-ON’ induction in the presence of cyanamide[40, 41].

We constructed a two-vector system to introduce the ^I-^*Sce*I expression cassette under the control of the *DDI2* promoter and test whether cyanamide-inducible ^I-^*Sce*I expression worked for selectable marker excision (**Figure 1A**: pITESceIcmd & pITEvd1). In this system, a 174 bp repeats were surrounding the ^I-^*Sce*I sites located at both sides of the hygromycin resistance selectable marker (*HphMX*). These two plasmids were co-transformed into CEN.PK113-32D [46] genome with the replacement of the *HO* gene, the gene required for mating-type switch in natural *S. cerevisiae* isolates. Colony PCR was used to confirm that the transformation in four transformants was successful (**Figure 1A**: strain oQIE0). To induce the ^I-^*Sce*I expression, we tried to use 4 mM cyanamide and 2 mM cyanamide, but the concentration at 4 mM slowed the growth in our strains. We grew oQIE0 in YPD broth supplemented with 2 mM cyanamide from a low OD600 ∼ 0.001 for 20 hours and then isolated single colonies by spreading the diluted culture on a YPD agar plate supplemented with 2 mM cyanamide. We tested 27 colonies through yeast colony PCR, and the Hph gene was successfully removed in 67% of the colonies, shown by the loss of the amplification of PCR fragment #3 **(Figure 1A**: generating strain oQIE1).

**Figure 1.**
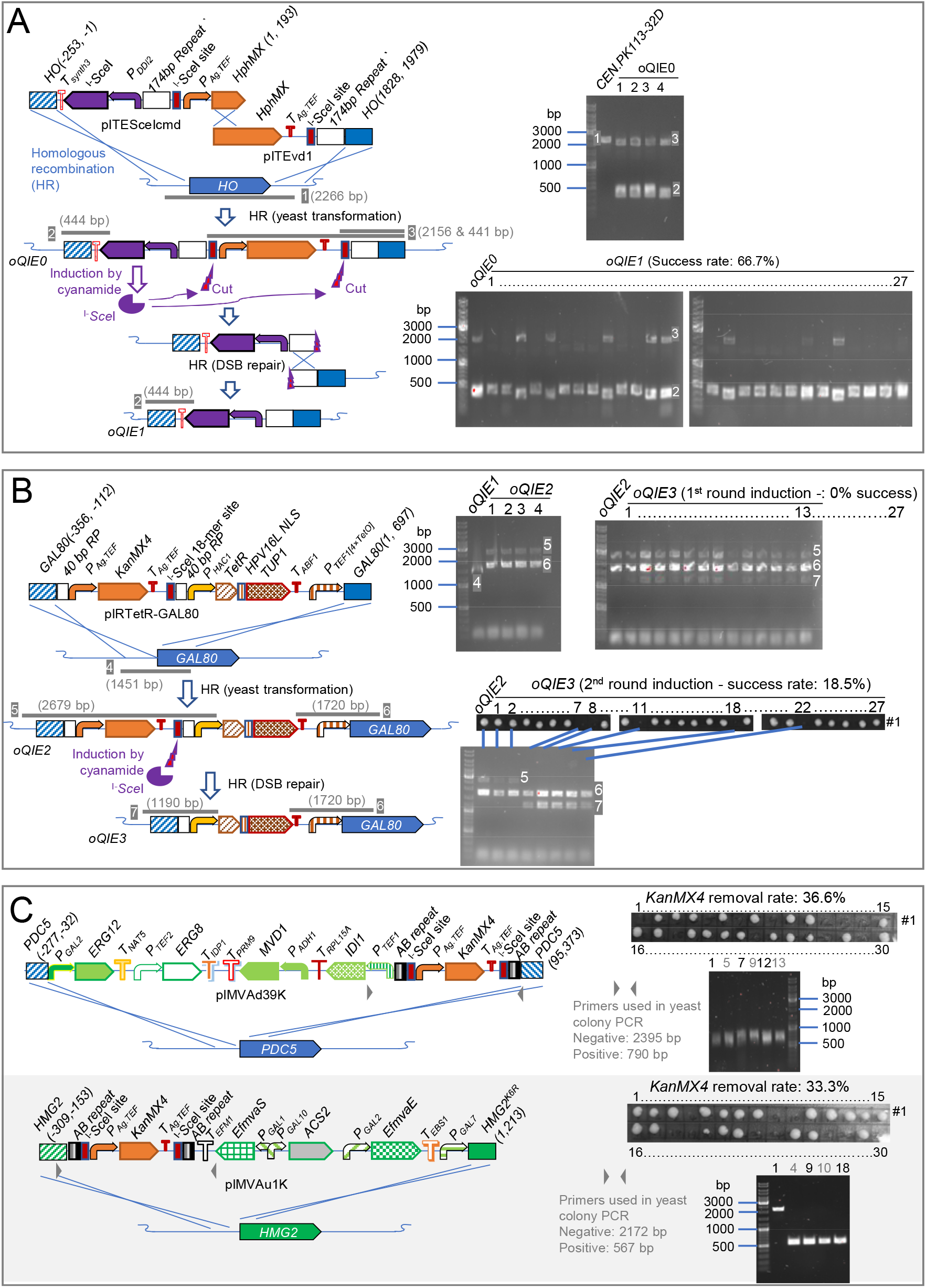
Cyanamide-inducible expression of ^I-^*Sce*I facilitating selectable marker removal for iterative strain engineering. (A) Integration of cyanamide-inducible ^I-^*Sce*I cassette at *ho* locus and selectable marker removal. (B) Integration of tetracycline-inducible *GAL80* module for tetracycline-mediated repression on galactose-inducible promoters and selectable marker removal. (C) Integration of the mevalonate pathway modules and selectable marker removal. Negative, the *KanMX4* marker being not removed; Positive, the *KanMX4* marker being removed. *P*_*XXX*_, the promoter of gene *XXX*; *T*_*XXX*_, the terminator of gene *XXX. T*_*synth3*_, a synthetic minimal terminator. NLS, nucleus-localising sequence. *TetR-HPV16L*^*NLS*^*-TUP1*, encoding a tetracycline-depressible repressor. TetO, TetR-binding sequence. Metabolic pathway genes for terpene precursor synthesis: *ERG12, ERG8, MVD1, IDI1, EfmvaS, ACS2*, and *EfmvaE. GAL*, the genes involving in galactose metabolism and regulation. #1, yeast clones replicated on the YPD agar supplemented with 300 μg ml^-1^ G418. The yeast colony PCR results are shown in DNA agarose gel images.

At the next step, we transformed the construct that encoded the genetic circuit for tetracycline-mediated repression of *GAL* promoters (**Figure 1B**: pIRTetR-GAL80) [38]. This construct is targeted to replace the promoter of the *GAL* repressor gene *GAL80* with the G418 selectable marker *KanMX4*, the expression cassette of an artificial transcription factor TetR-Tup1p, and the *TEF1* promoter inserted with 4 TetO elements. This construct allows tetracycline-mediated induction of the *GAL80* gene, as the result, *GAL* promoters are repressed in the presence of tetracycline and de-repressed in the absence of tetracycline. In this construct, we have previously cloned a single ^I-^*Sce*I site at the downstream of G418 resistance marker gene *KanMX4* and two 40 bp homologous repeats at the both ends of *KanMX4-*^*I-*^*SceI* site sequence. This construct was successfully transformed into strain oQIE1 to generate strain oQIE2 and successful transformants were confirmed by yeast colony PCR. We performed cyanamide induction and characterised 27 colonies through yeast colony PCR. However, none of these colonies showed a successful and complete removal of the *KanMX4* marker. Instead, the colony PCR results showed that there were two genotypes: *KanMX4* unremoved (**Figure 1B:** PCR fragment #5) and *KanMX* removed (**Figure 1B:** PCR fragment #7). We chose first four colonies to perform the second round of cyanamide induction and tested 27 colonies for their resistance against G418. Five colonies had the *KanMX* marker removed, which generated strain oQIE3.

In further strain engineering, we aimed to re-construct the yeast strain harbouring the augmented mevalonate pathway, which required the transformation of two plasmids pIMVAd39T (Addgene #98303) and pIMVAu1 (Addgene #98301)[47]. In these two plasmids, autotrophic selectable markers were used for yeast transformation, but were not available in oQIE3. We further used the 80-mer AB repeats, published in the selectable marker cassette project[28], to mediate homologous recombination-based repair of ^I-^*Sce*I-induced DSBs. We replaced the selectable marker with *AB-*^*I-*^*SceI-KanMX4-*^*I-*^*SceI-AB* fragment in pIMVAd39T and pIMVAu1, and sequentially transformed them into oQIE3 with cyanamide-induced marker removal (**Figure 1C**). In both cases, the efficiency of marker removal was not as high as that when galactose-inducible promoter was used for ^*I-*^*SceI* induction. The marker removal was successful in these two rounds of strain engineering with a success rate ≥ 33%.

### *In situ* gene replacement design for plasmid-free cloning of heterologous promoters into yeast genome

The fundamental principles in metabolic engineering and synthetic biology rely on the expression of heterologous genes under the control of different promoters [48]. These promoters can be either constitutive or regulatory and show a range of transcription strengths[49]. Nevertheless, promoter characterisation is essential to reveal their fundamental genetic features in host organisms. These genetic features are then referred to as engineering dictionary contents for rational design of genetic regulatory networks for metabolic control or other applicational aims. In *S. cerevisiae*, promoters can be characterised using an episomal plasmid system or a genome-integrating system. Genome-integrating systems are ideal to deliver a genetically homogeneous cell population and allow for characterisation independent from the requirements of specific selection pressure for plasmid maintenance. When using genome-integrating system to characterise yeast promoters, the classical molecular cloning procedure is followed, including cloning yeast promoters into an *E*.*coli*/*S. cerevisiae* shuttle plasmid and transforming the resultant plasmid into *S. cerevisiae*. This procedure is time-consuming and labour-intensive, including the steps of PCR preparation of promoter fragments, preparation of vector fragment, fragment-vector assembly/ligation, *E. coli* transformation, PCR to screen for positive clones, plasmid extraction, plasmid linearisation, yeast transformation, yeast clone screening and verification, and then promoter characterisation. We aimed to shorten this procedure into four essential steps: PCR preparation of promoter fragments, yeast transformation, yeast clone screen and verification, and then promoter characterisation. Applying cyanamide-inducible ^I-^*Sce*I expression, we designed an *in situ* gene replacement system that allows direct genomic integration of promoter fragments at the upstream of a reporter gene (**Figure 2**).

**Figure 2.**
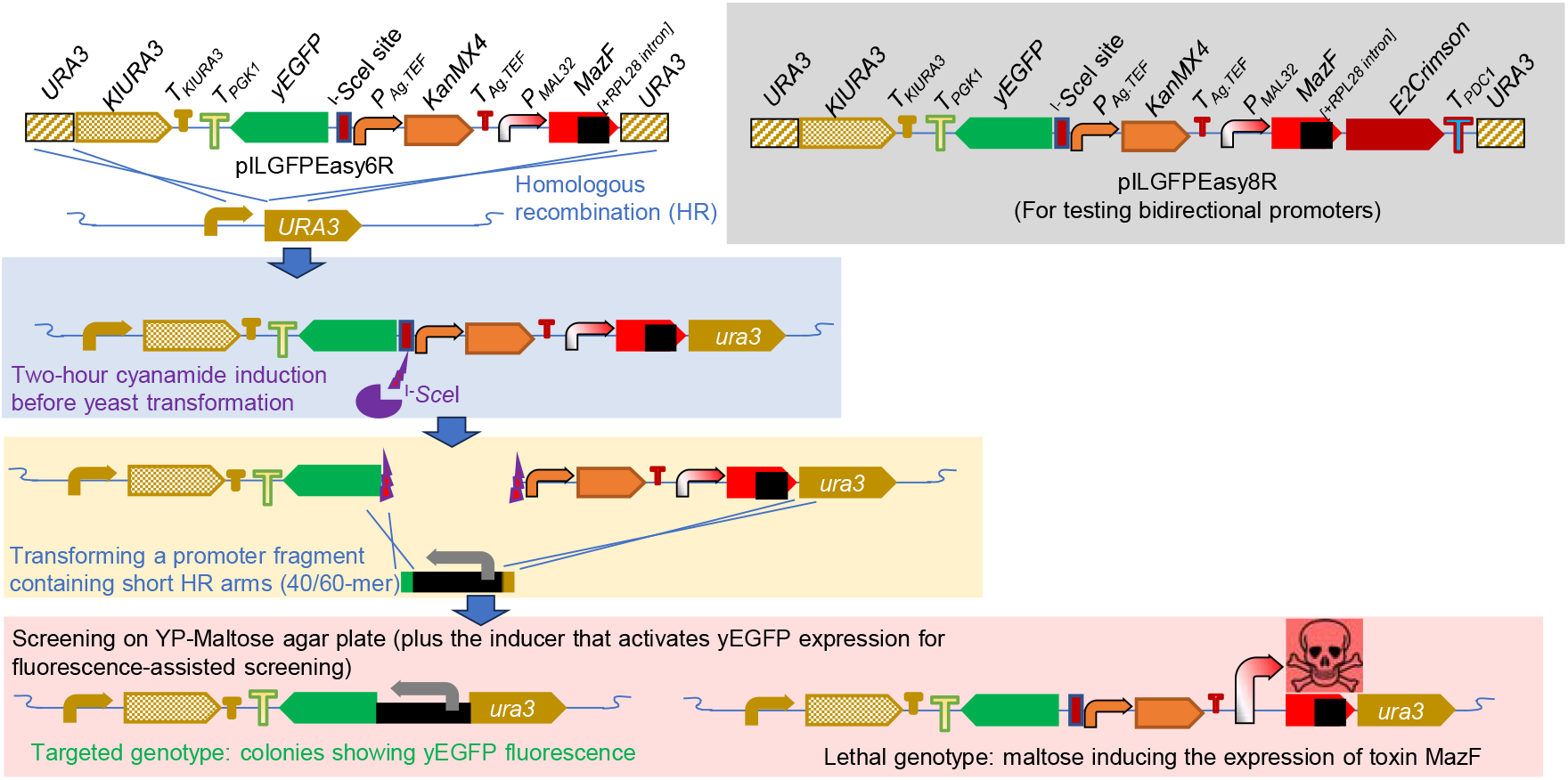
Diagrams of *in situ* gene replacement design for plasmid-free cloning of heterologous promoters into yeast genome. *P*_*XXX*_, the promoter of gene *XXX*; *T*_*XXX*_, the terminator of gene *XXX*. MazF, encoding an *E. col*i interferase toxin.

To test this design, we developed two plasmids, pILGFPEasy6R and pILGFPEasy8R. Both plasmids can be integrated into the yeast genome at the *ura3* locus, containing a *KlURA3* selectable marker for transformation selection in ura3 background strain, *yEGFP* gene fused with the *PGK*1 terminator, an ^I-^ *Sce*I site, a G418-resistant selectable marker *KanMX4*, a *MAL32* promoter-controlled toxin gene *MazF* inserted with the *RPL28* intron. In plasmid pILGFPEasy8R, a E2Crimson reporter fused with the *PDC1* terminator is inserted at the downstream of MazF and in the opposite direction of yEGFP gene.

In theory, either pILGFPEasy6R or pILGFPEasy8R fragments can be transformed into a background strain harbouring a cyanamide-inducible ^I-^*Sce*I expression cassette. In the resulting strain of pILGFPEasy6/8R transformation, cyanamide is added two hours before yeast transformation to induce ^I-^*Sce*I expression. During the yeast transformation, ^I-^*Sce*I-mediated DSBs are repaired through homologous recombination by the promoter PCR fragment with short homology arms. The transformation cells are precultured to allow sufficient cell division to generate cells with the MazF expression cassette removed. The cell mixtures were spread onto YP-maltose agar, if necessary, supplemented with the inducers to activate the heterologous promoter and drive yEGFP expression. As the MazF toxin is induced by maltose, only the cells with the MazF expression cassette removed can form colonies.

### Characterisation of regulatory promoters in *S. cerevisiae*

To test the cloning procedure of *in situ* gene replacement, we aimed to characterise a set of regulatory promoters, which were regulated by either endogenous regulatory mechanisms or synthetic regulatory mechanisms. We constructed a yeast strain (strain oQR36o). In strain oQR36o, four heterologous expression cassettes were introduced at the *HO* locus: (1) an artificial transcription factor (TF) Zif268-hER-VP16^A^ (the fusion of Zif268 zinc finger DNA binding domain, estrogen sensing domain, and VP16 transactivating domain) was expressed under the control of the YPK2 promoter, a weak promoter; (2) an artificial TF LmrA-SV40^NSL^-Mig1^C^ (the fusion of a bacterial TetR-like TF LmrA, SV40 nucleus-localising sequence, and Mig1p C-terminal trans-repression domain) was expressed under the control of the *ADR1* promoter; (3) an artificial TF TetR-HPV16L^NLS^-Tup1 (the fusion of tetracycline repressor TetR-HPV16L nucleus-localising sequence, and trans-repressing TF Tup1) was expressed under the control of the *HAC1* promoter; and (4) ^I-^*Sce*I was expressed under the control of cyanamide-inducible *DDI2* promoter.

Plasmid pILGFPEasy6R was transformed into strain oQR36o for characterisation of mono-directional promoters, and plasmid pILGFPEasy8R was transformed for characterisation of bidirectional promoters. Noted is that pILGFPEasy8R covers pILGFPEasy6R function. Each promoter fragment was amplified through PCR, including 40 or 60-bp homologous recombination arms for genomic integration. Yeast transformation was performed using the PEG/LiAc/ssDNA protocol with a two-hour cyanamide induction before transformation. After heat shock, cells were collected by centrifuging, resuspended in 1 ml YPD, and left at room temperature overnight. In the next day, cells were mixed and spread to agar plates. At this step, it is recommended to spread 10 μl or less cells per plate, otherwise it is difficult to isolate single colonies. After 48-hour incubation at 30 °C, the plates were imaged using a BioRad ChemiDoc Imaging System. Although brighter colonies were only a small proportion of those plated, they were found to be correctly transformed when selected (**Supplementary Figure 1**).

We further constructed another plasmid (pILGFPEasy9R; **Supplementary Figure 2**) by inserting the *P*_*DDI2*_*-*^*I-*^*SceI-T*_*synth3*_ expression cassette into the downstream of the *PDC1* terminator in plasmid pILGFPEasy8R. The pILGFPEasy9R construct was transformed into strain CEN.PK113-5D. Using the resulting strain, we transformed *S. kudriavzevii GAL1* promoter [50] to test the efficiency of *in situ* gene replacement. Consistently, brighter colonies were a smaller population, but they were found to be correctly transformed when selected.

In total, we characterised five groups of inducible promoters. They are (1) tetracycline-inducible *TEF1[4×TetO]* promoters, (2) cyanamide-inducible *DDI2* promoter, (3) synthetic β-estradiol-inducible promoters, (4) maltose-inducible *MAL32/31* bidirectional promoters, and (5) five galactose-inducible *GAL10/1* promoters.

We previously synthesised a *TEF1[4×TetO]* promoter by inserting four *TetO* motifs between the core promoter and upstream activation sequence (UAS) of the *TEF1* promoter [38]. The *TEF1[4×TetO]* promoter is repressed in the strain expressing TF TetR-HPV16L^NLS^-Tup1 (TetR-Tup1) and is inducible by tetracycline addition. The originally tested *TEF1[4×TetO]* promoter (Figure 3A: UAS-579), containing a full-length UAS, demonstrates a moderate transcription strength; half of the strength of the unmodified *TEF1* promoter. The tetracycline-inducible promoters with weaker observed strength might be useful for controlling certain expression of certain genes, like the TF-encoding gene. We hypothesised that the weaker promoters could be developed through truncating UAS sequences. We then tested two truncated *TEF1[4×TetO]* promoters, UAS-382 (the promoter starting at -382 position of the *TEF1* gene) and UAS-288 (the promoter starting at -288 position). Consistent with the hypothesis, shorter *TEF1[4×TetO]* promoter showed weaker transcription.

**Figure 3.**
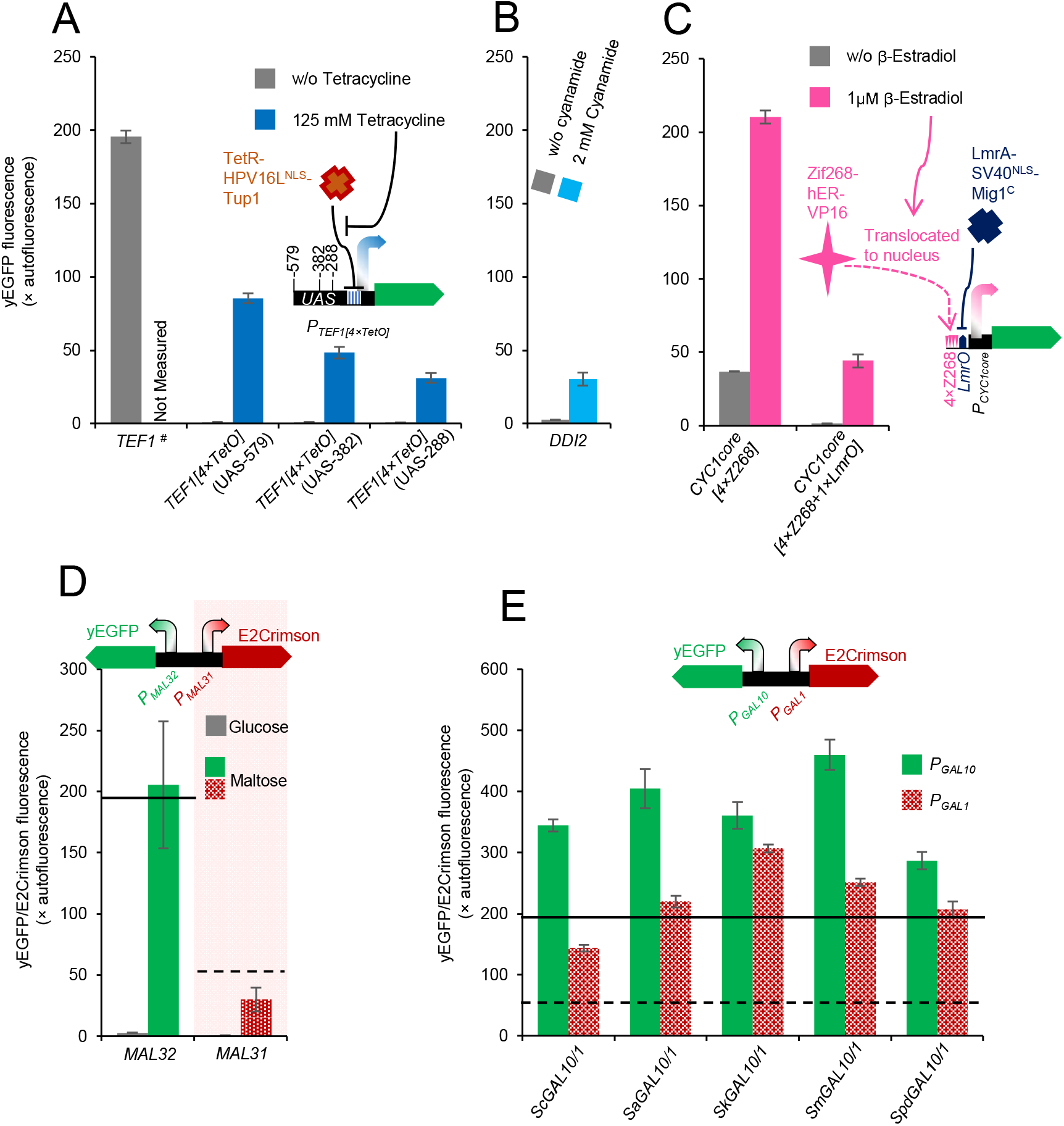
Characterisation of regulatory promoters in *S. cerevisiae*. (A) Tetracycline-inducible *TEF1[4×TetO]* promoters: truncating the upstream activation sequence (UAS) resulted in decreased expression outputs. TetR-HPV16L^NLS^-Tup1, an artificial trans-repressor binds to TetO sequences. Tetracycline binding to TetR disassociates the TetR-TetO interaction. (B) Cyanamide-inducible *DDI2* promoter. (3) Estrogen (β-estrodiol)-inducible promoters. Zif268-hER-VP16, an artificial trans-activator targeting to Z268 sequences. β-Estrodiol binding to hER (human estrogen receptor) resulting in nuclear translocation. LmrA-SV40^NSL^-Mig1^C^, an artificial trans-repressor binding to the *LmrO* sequence. (D) Maltose-inducible promoters. (E) Galactose-inducible promoters. The yEGFP fluorescence under the control of the *TEF1* promoter characterised on glucose are represented as bold lines and the E2Crimson fluorescence under the control of the *TEF1* promoter characterised on glucose are represented as bold lines in (D & E). Cells were grown in YNB-glucose media (A and C), YP-glucose media (B & D), YP-maltose media (D), or YNB-galactose media (E). Mean values ± standard deviation were shown (n =3 biological replicates).

We characterised the *DDI2* promoter which had been used to control ^I-^*Sce*I expression in this study. In previous studies [40, 41], the *DDI2* promoter was not characterised with an external reference. Our tests showed that the strength of the *DDI2* promoter was ∼6.4-fold lower than the *TEF1* promoter (**Figure 3B**). This may explain the cause of the lower marker removal efficiency when ^I-^*Sce*I was under the control of the *DDI2* promoter in comparison to that when the strong *GAL1* promoter was used to control ^I-^*Sce*I expression.

Synthetic estrogen-inducible promoters were used in early synthetic biology studies [51]. However, these promoters were leaky, i.e., for the *CYC1* core promoter fused with Zif268-binding elements (**Figure 3C**) [38] and the *GAL1* promoter inserted with Zif268-binding elements[39]. To solve this problem, we hypothesized that an artificial trans-repressor could be used to repress its basal expression. Here, we tested a synthetic *CYC1*core promoter fused with four Z268 (Zif268-binding) elements and one LmrA-binding LmrO sequence (*CYC1core[4×Z268+1×LmrO]*). The *CYC1core[4×Z268+1×LmrO]* showed a very low basal expression, and was inducible by estrogen β-estradiol, but the activity was ∼4.7-fold lower than the *CYC1core[4×Z268]* promoter (Figure 3C).

For bidirectional promoters, we characterised maltose-inducible *MAL32/31* promoters and galactose-inducible *GAL1/10* promoters from five *Saccharomyces* species: *S. cerevisiae* (*ScGAL10/1*), *S. arboricola* (*SaGAL1/10*), *S. kudriavzevii* (*SkGAL1/10*), *S. mikatae* (*SmGAL1/10*), and *S. paradoxus* (*SpdGAL1/10*). The *MAL32/31* and Sc*GAL1/10* promoters have been have been characterised for bidirectional transcriptional activation and strength previously [50]. For the *SaGAL1/10, SkGAL1/10, SmGAL1/10*, and *SpdGAL1/10* promoters, we previously measured the strength at the *GAL1* direction, and the data for *GAL10* directions were not available to guide their applications in metabolic engineering. We therefore used our system to re-characterise their activities. As the external reference for E2Crimson measurement, the strength of the *TEF1* promoter was characterised on glucose using E2Crimson, which was under the control of the *PGK1* terminator (Figure 3D & 3E: dashed line). Consistent with the previous study, both sides of the *MAL32/31* promoter were induced by maltose. The *MAL32* promoter was induced to the similar level of the *TEF1* promoter, and the *MAL31* promoter was at the halved level of the *TEF1* promoter.

The *GAL1/GAL10* promoters were characterised on galactose in a *GAL80* wildtype background. These conditions were different from our previous tests in *gal80*Δ background in the whole batch cultivation with glucose as the initial carbon source. Here, we primarily investigated the potential maximal strength of these *GAL1/GAL10* promoters under galactose induction. Using E2Crimson as the reporter, the output from the *ScGAL1* promoter was ∼2.6-fold of that from the *TEF1* promoter. This fold-change was in the same range with the ∼2.5 fold in our previous measurement, when yEGFP was used as the reporter. This indicates that our tests were consistent although some experimental conditions were changed. For the *GAL1* promoters, the strength was sequenced as: *SkGAL1* > *SmGAL1* > *SaGAL1/SpdGAL1* > *ScGAL1*. Fortunately, all five *GAL10* promoters were stronger than the *TEF1* promoter. The strongest *GAL10* promoter was the *SkGAL10* promoter, showing the output ∼2.4-fold higher than that of the *TEF1* promoter. The *SpdGAL1* promoter was the weakest *GAL10* promoter, ∼1.5-fold of the *TEF1* promoter. These suggest that these five bidirectional *GAL1/10* promoters are good options for gene overexpression in metabolic engineering.

## Discussion

In this study, we tested two yeast engineering methods: cyanamide-induced ^I-^*Sce*I-mediated double-strand DNA breaks (DSBs) for selectable marker removal and the plasmid-free *in situ* promoter replacement through cyanamide-induced ^I-^*Sce*I-mediated DSB and maltose-induced MazF-mediated negative selection. Although not perfect in the success rates for target engineering, both methods can practically facilitate routine yeast engineering. Applying cyanamide-mediated gene induction allows its compatibility with pathway engineering using the galactose-inducible promoters. The *in situ* promoter replacement system minimises the operational steps for characterisation of yeast promoters through genome integration. Both systems can be further improved and may be replicable in other non-conventional yeast platforms.

The cyanamide-inducible *DDI2* promoter was used in the aim to deliver orthogonal control on different genetic engineering tools. Apparently, both engineering methods apply the *DDI2* promoter and ^I-^*Sce*I meganuclease in this study. This may cause compatibility problems when they were applied simultaneously in the same cell. In further development, alternative small-molecule-inducible genetic circuits can be considered, i.e., the ligand/factor pairs of 2,4-diacetylphloroglucinol/PhlF[52], camphor/CamR[52], 1,2-bis(4-hydroxyphenyl)ethane-1,2-dione/hER mutant[53], steroids/mammalian type I nuclear receptors[54]. These inducible circuits may not only be useful to control alternative meganucleases or genome-engineering tools, but also alternative killing toxins (like MazF; **Figure 3**) for negative selections.

Nevertheless, it might be necessary to optimise the regulatory promoters for a tight ON/OFF response and minimise the promoter lengths [55] to facilitate PCR-based cloning and assembly, which can be facilitated by the plasmid-free *in situ* promoter replacement. We have previously identified that the fusion of 4×Z268-containing DNA sequence to the *CYC1* core promoter contributed to the basal expression [38], possibly due to endogenous *trans*-activators activating the transcription. Here, we solved the leakiness problem in a β-estradiol-inducible genetic circuit by introduction of a synthetic *trans*-repression mechanism in the regulatory promoter (**Figure 3C**). The high affinity of a synthetic *trans*-activator to its target binding sites might also result in the leaky *trans*-activation[56]. To solve this problem, using a low-affinity DNA-binding domain eliminated the leakiness at the expense of the induction strength. To increase the induction output strength in this case, a molecular clamp was used to stabilise the interaction of multiple synthetic *trans*-activators and target DNA motifs at the target promoter. In the tetracycline-inducible circuit case, we showed that the length of upstream activation sequence (UAS) determined the induction strength (**Figure 3A**). Consistently, tandem assembly of multi-UASs has been used to engineer super strong yeast promoters[57]. These emphasise that there are alternative solutions in synthetic biology to achieve similar biological effects and genetic circuit design principles shall be discovered for precise genetic design.

In summary, we prototyped the cyanamide-induced meganuclease tools to facilitate *in situ* genome engineering in *S. cerevisiae*. These tools may facilitate strain engineering to establish genetic engineering at multiple genome loci and the characterisation of yeast promoters. Availability of multiple fine small-molecule-inducible genetic circuits are important to develop more complexed networks for precise control on more genome engineering tools in a single cell [58, 59]. In this scenario, it will be also important to develop computer software so that individual genome-engineering tools may work synergistically in a biofoundry setting to power the engineering biology capacity for synthetic biology and metabolic engineering applications [60].

## Acknowledgements

This research was supported by the Australian Government through the Australian Research Council Centres of Excellence funding scheme (project CE200100029). The views expressed herein are those of the authors and are not necessarily those of the Australian Government or Australian Research Council.

