## Supplementary Data 1 for "Cyanamide-inducible expression of homing nuclease ^I-^*Sce*I for iterative genome engineering and parallel promoter characterisation in *Saccharomyces cerevisiae*"

Supplementary Figure 1.

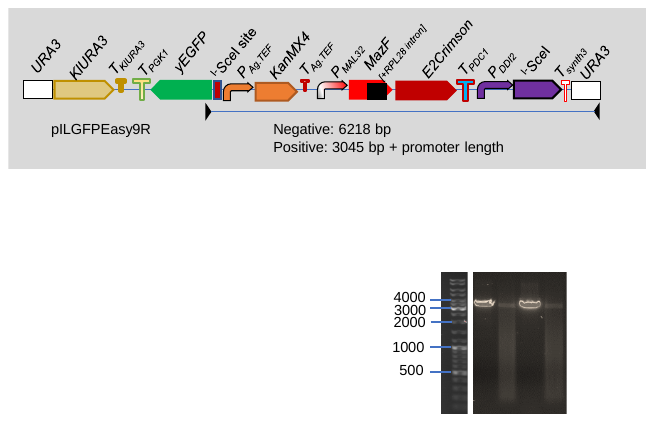

Supplementary Figure 2.
